# A single factor for safer cellular rejuvenation

**DOI:** 10.1101/2025.06.05.657370

**Authors:** Lucas Paulo de Lima Camillo, Rihab Gam, Katsiaryna Maskalenka, Francis J. A. LeBlanc, Gustavo Antonio Urrutia, Gabriel M. Mejia, Henry E. Miller, Christopher P. Wardlaw, Adam Pickles, Laura Everton, Ringaile Zaksauskaite, Rejina B. Khan, Andreas Welsh, Samira Gambo, Stephany Gallardo, Zoryana Oliynyk, Sagar S. Varankar, Alexander E. Epstein, Adam Bendall, Jonathan Mowatt, Daniel Ives, Brendan M. Swain

## Abstract

Ageing is a key driver of the major diseases afflicting the modern world. Slowing or reversing the ageing process would therefore drive significant and broad benefits to human health. Previously, the Yamanaka factors (OCT4, SOX2, KLF4, with or without c-MYC: OSK(M)) have been shown to rejuvenate cells based on accurate predictors of age known as epigenetic clocks. Unfortunately, OSK(M) induces dangerous pluripotency pathways, making it unsuitable for therapeutic use. To overcome this therapeutic barrier, we screened for novel factors by optimising directly for age reversal rather than for pluripotency. We trained a transcriptomic ageing clock, unhindered by the low throughput of bulk DNA methylation assays, to enable a screen of unprecedented scale and granularity. Our platform identified SB000, the first single gene intervention to rejuvenate cells from multiple germ layers with efficacy rivalling the Yamanaka factors. Cells rejuvenated by SB000 retain their somatic identity, without evidence of pluripotency or loss of function. These results reveal that decoupling pluripotency from cell rejuvenation does not remove the ability to rejuvenate multiple cell types. This discovery paves the way for cell rejuvenation therapeutics that can be broadly applied across age-driven diseases.

**Highlights:**

- SB000 drives multi-omic rejuvenation in human fibroblasts, as evidenced by substantial reversal of numerous epigenetic clocks, lowered single-cell transcriptomic age, and decreased senescence-associated gene expression.
- In contrast to OSK(M), SB000 treatment maintains transcriptomic and functional measures of fibroblast identity without the activation of pluripotency.
- SB000 rejuvenation generalises to keratinocytes, cells from another germ layer, with potency matching or surpassing OSK(M).

## Introduction

Ageing involves a gradual decline in biological function that contributes to a wide range of human diseases, including cardiovascular disease, neurodegeneration and osteoarthritis [1]. For millennia, this fact has motivated humanity to search for solutions, with ancient scholars referencing the search for a rejuvenating elixir as far back as the 5th Century BCE.

In modern times, the search has yielded several interventions that slow ageing and extend lifespan across model organisms. Rapamycin, an allosteric inhibitor of the mechanistic target of rapamycin (mTOR), extends median lifespan by up to 26% in mice and exerts similar benefits in yeast, worms, and flies [2, 3, 4, 5]. Caloric restriction (CR) without malnutrition remains the best-known geroprotector, increasing lifespan by up to 60% in rodents, with efficacy in yeast, worms, flies, mosquitoes, killifish, and dogs [6, 7, 8, 9, 10, 11, 12]. However, despite their potency, both rapamycin and CR primarily seem to slow the accrual of damage during ageing rather than reverse it. When mice return to an ad libitum diet after CR, they rapidly reacquire baseline mortality risk [13]. Likewise, transient post-development rapamycin treatment confers mixed effects on longevity [14, 15, 16]. Overall, the current pharmacological arsenal largely comprises agents that decelerate rather than rewind biological time.

A striking exception is the Yamanaka factors, a quartet of transcription factors OCT4, SOX2, KLF4, and c-MYC (OSKM). Induced expression of OSKM converts differentiated somatic cells into induced pluripotent stem cells (iPSCs), a discovery worthy of the 2012 Nobel Prize [17]. More recently, these factors have garnered interest from the ageing research field due to a putative side effect: rejuvenation. In 2013, Steve Horvath applied his multi-tissue DNA methylation clock to iPSCs, noting that their predicted age was near or below zero [18]. In 2019, a post-hoc analysis of an *in vitro* OSKM timecourse showed a remarkable decrease of several decades in the predicted age from multiple epigenetic clocks over the course of a few weeks [19], which can be induced even in cells from super-centenarians [20].

The Yamanaka factors also have systemic effects *in vivo*. In a seminal study, Ocampo *et al.* showed that cyclic OSKM expression can extend the lifespan of progeroid mice by around 33% [21], subsequent work revealing that this treatment also improves the health of aged wild-type mice [22]. Since then, the three-factor cocktail OSK extended the remaining lifespan of aged wild-type mice when expression was limited to senescent cells [23] or after systemic delivery with adeno-associated virus (AAV) [24]. OSK induction has also been linked to short-term disease amelioration in young mice, suggesting that it can impart resilience in an age-independent manner [25]. Today, the Yamanaka factors are the gold standard for cellular rejuvenation.

The therapeutic potential of OSK(M) for human therapy is, however, limited by safety concerns given the narrow therapeutic window. Continuous OSKM expression in mice causes rapid death due to widespread loss of cell identity, primarily from liver and intestinal failure [26, 21]. Shorter protocols often produce teratomas, tumours containing tissues from all germ layers, akin to a body without a body plan [27, 28, 29]. To avoid progression to pluripotency, many have focused on transient expression of the Yamanaka factors, or similar gene sets. Cells undergoing transient reprogramming with OSKM appear to revert to their original cell identity after the factors’ expression stops, while maintaining a reduced epigenetic age [30]. However, the safe dosing and timing window is narrow, even in well-controlled *in vitro* experiments, leading many groups to remove c-MYC from the cocktail to avoid its oncogenic effects. Unfortunately, the OSK trio of genes is also able to form pluripotent colonies *in vitro* [31]. As even a single pluripotent cell in a patient could be enough to form a tumour, there are reservations about OSK(M)’s development as a broad rejuvenation therapeutic.

While the Yamanaka factors are a remarkable tool, they were discovered through the optimisation of pluripotency. A search for generalisable and safe interventions using a “rejuvenation first” approach is warranted. Rigorous optimisation of rejuvenation therapies requires sensitive, high-throughput biomarkers of biological age. Unfortunately, the search has been hampered by a lack of suitable tools for highly parallelised rejuvenation screens. Epigenetic clocks, statistical models that predict age from DNA methylation states, are the most accurate such biomarkers to date but typically rely on low-throughput bulk assays requiring large populations of cells [32]. We therefore constructed what is, to our knowledge, the most precise single-cell transcriptomic ageing clock in human dermal fibroblasts. Ageing Clock 3 (AC3) enables perturbation screens at scale with thousands of multiplexed genetic interventions, allowing us to identify candidate rejuvenation factors.

Here we highlight SB000, a single gene uncovered by AC3 that rivals OSKM’s rejuvenating potency and exceeds OSK’s safety. In human dermal fibroblasts, SB000 lowers single-cell transcriptomic age according to AC3 and attenuates harmful transcriptomic signatures of multiple hallmarks of ageing, including loss of proteostasis, increased sterile inflammation, and dysregulated intercellular communication, amongst others. Moreover, SB000 reduces DNA methylation age across more than a dozen epigenetic clocks. Crucially, fibroblast identity markers such as collagen secretion are preserved without induction of pluripotency, as shown by the absence of pluripotent colonies in long-term culture, in stark contrast to OSK. The efficacy of SB000 also generalises beyond mesoderm-derived dermal fibroblasts: it rejuvenates lung fibroblasts and, notably, ectoderm-derived keratinocytes, indicating broad applicability across germ-layer boundaries.

SB000 is the first single rejuvenation gene discovered *de novo* through high-throughput single-cell screening tailored for safe, multi-cell-type rejuvenation. Our findings showcase how data-driven design, coupled with quantitative ageing clocks, can accelerate the discovery of next-generation geroprotective interventions with the potential to significantly extend human healthspan and lifespan.

## Results

### Using a transcriptomic clock to discover rejuvenation interventions

Accurately quantifying biological age at the level of individual cells accelerates the systematic screening of rejuvenation therapeutics by several orders of magnitude. We therefore trained a transcriptome-based ageing clock, unhindered by the low throughput of bulk DNA methylation (DNAm) assays, to enable screens of unprecedented scale and granularity. We assembled a primary human dermal fibroblast (HDF) compendium from more than 100 donors spanning 1 to 87 years of age, comprising male and female samples drawn from Caucasian, Black, Hispanic, and Arabic ancestries, establishing the AC3 training cohort (Figure 1A).

**Figure 1:**
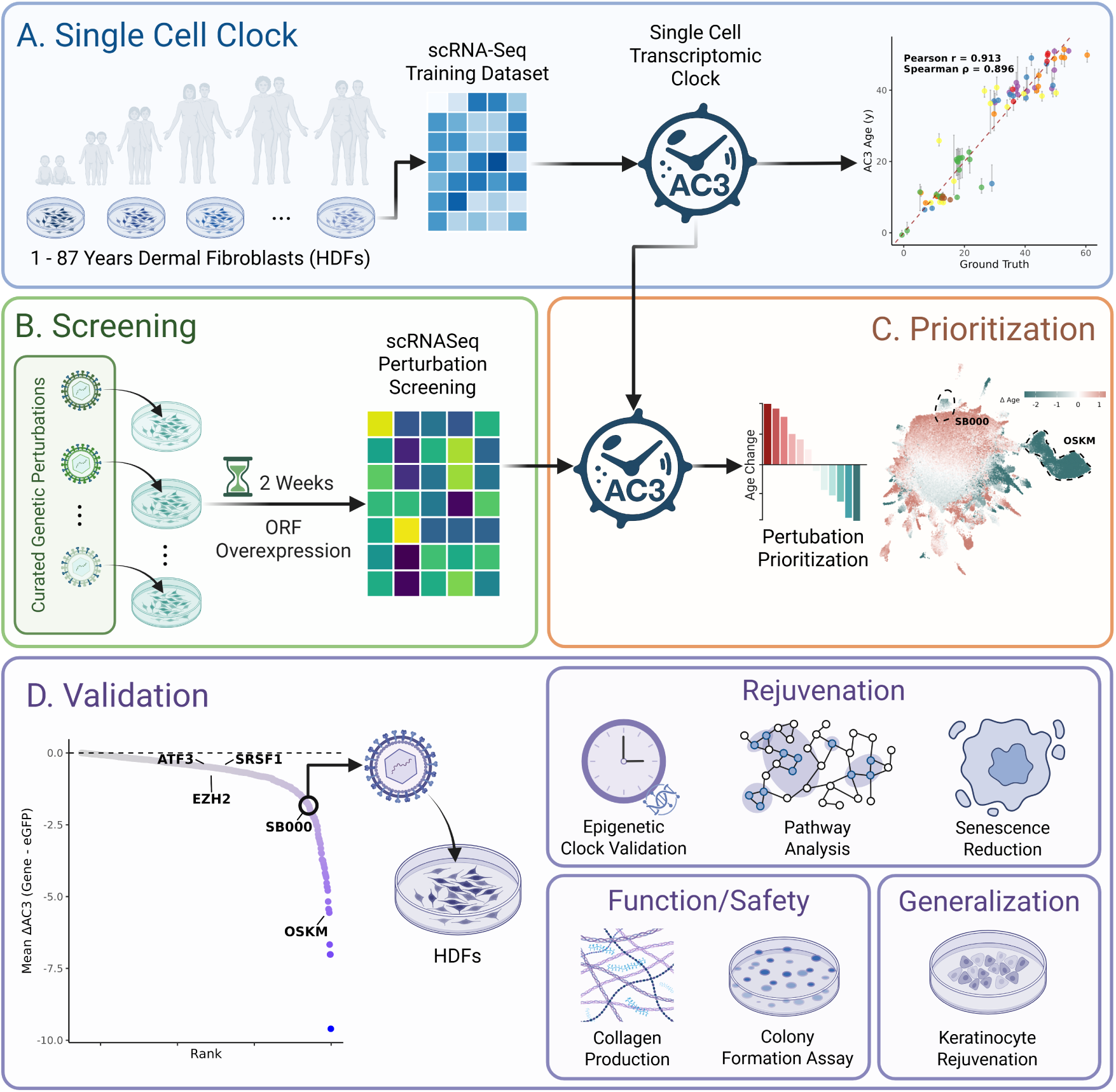
Using a transcriptomic clock to discover rejuvenation interventions. **(A)** Shift’s discovery platform relies on a large dataset of primary aged cells, on which the single-cell transcriptomic age predictor AC3 was trained. AC3 predicts the age of single cells with state-of-the-art accuracy. The predicted age of cells is shown on the y-axis vs. the ground truth biological age according to the PCSkinAndBlood clock on the x axis. Points and error bars represent median *±* IQR for each sample, coloured by batch in the test dataset. Inset is the Pearson’s correlation coefficient on the single-cell level. **(B)** AC3 allows large single-cell screens, in which the single-cell transcriptome of cells is captured after transduction with candidate rejuvenation genes. **(C)** An example screen, where 1500 genes were screened for an effect on cellular age using AC3 *in vitro*. Shown is a UMAP projection of cells in this dataset, coloured by the change in AC3-predicted age for each cell relative to the eGFP control for each donor. **(D)** Genes were then ranked based on the mean reduction in AC3-predicted age; SB000 and selected comparator genes from the literature are labelled. We then performed further characterisation experiments with SB000 for efficacy, safety and generalisation.

In order to construct and validate AC3, we applied our proprietary machine learning algorithm to the single-cell transcriptomes of the HDF cohort and benchmarked the resulting age predictions against the sample’s DNAm age according to established epigenetic clocks. Across a large independent test dataset containing both physiologically-aged and perturbed HDFs, AC3 achieved a single-cell Pearson correlation of 0.913 with the PCSkinAndBlood predictor [33] (Figure 1A). The performance rose to 0.946 when the median prediction for each sample was used to calculate the Pearson metric. The accuracy was reproducible across different single-cell chemistries (10X genomics, Scale Biosciences, and Parse Biosciences) and sequencing instruments (Illumina NovaSeq 6000 and Illumina NovaSeq X; data not shown). Remarkably, AC3 attained state-of-the-art accuracies without resorting to metacell aggregation or smoothing, in contrast to existing frameworks [34, 35, 36, 37], such that every cell in a multiplexed screen can be considered an individual experiment. These properties establish AC3 as, to our knowledge, the first genuinely single-cell ageing clock, combining high precision with a throughput sufficient for discovery at scale.

Leveraging AC3’s resolution, we generated a perturbation atlas by overexpressing 1,500 manually curated open-reading frames (ORFs) in primary HDFs from three aged donors, including genes previously implicated in aging and rejuvenation (Figure 1B). After two weeks of constitutive expression by lentivirus, we profiled the cells with single-cell RNA-sequencing (scRNA-seq, Figure 1C), computed AC3-predicted ages for every cell, and ranked the mean effect of each intervention relative to an enhanced green fluorescent protein (eGFP) control for each donor. The majority of genes exerted minimal influence on transcriptomic age (Figure 1D). The genes SRSF1, ATF3, and EZH2, which have been shown previously to ameliorate hallmarks of ageing *in vitro*, showed only a modest reduction [38, 39, 40]. However, a number of candidates produced a pronounced reversal, with several single genes matching or exceeding the efficacy of OSKM. These results demonstrate that coupling a high-accuracy single-cell ageing clock with a curated genetic library enables high-throughput identification of rejuvenation genes in HDFs.

### SB000 rejuvenates primary human fibroblasts

Following the perturbation atlas, we further characterised the rejuvenative capacity of a single exemplar among the top 50 candidates, hereafter designated SB000. We first sought to test whether the expression of SB000 would reverse multiple hallmarks of ageing in aged primary fibroblasts from skin (n=6: 1 male, 2 female, datasets HDF1 and HDF2) and lung (n=4: 3 male, 1 female, dataset HLF1) with efficacy rivalling the Yamanaka factors. We therefore compared SB000 with full (OSKM) and partial (OSK, OS, OK, O, S) Yamanaka factor cocktails across single-cell transcriptomes and bulk DNA methylomes.

In order to quantify the effect of SB000 on transcriptomic age, we transduced primary HDFs and human lung fibroblasts (HLFs) from aged donors with a lentiviral vector driving constitutive SB000 expression and profiled the cells with scRNA-seq after two weeks (Figures 2A to 2C). SB000 reduced the mean AC3 age of fibroblasts by 4.52 years (s.d. 2.98 years, p = 0.00364), demonstrating robust rejuvenation across independent mesoderm-derived cell types (Figure 2D). Across fibroblasts of both organs, SB000 displayed a potency comparable to the full Yamanaka factor cocktail OSKM (5.46 years mean, 1.65 years s.d., p = 0.0290), supporting the results observed in the initial perturbation atlas. These data support that SB000 drives substantial rejuvenation in the transcriptomes of fibroblasts from both dermal and lung lineages.

**Figure 2:**
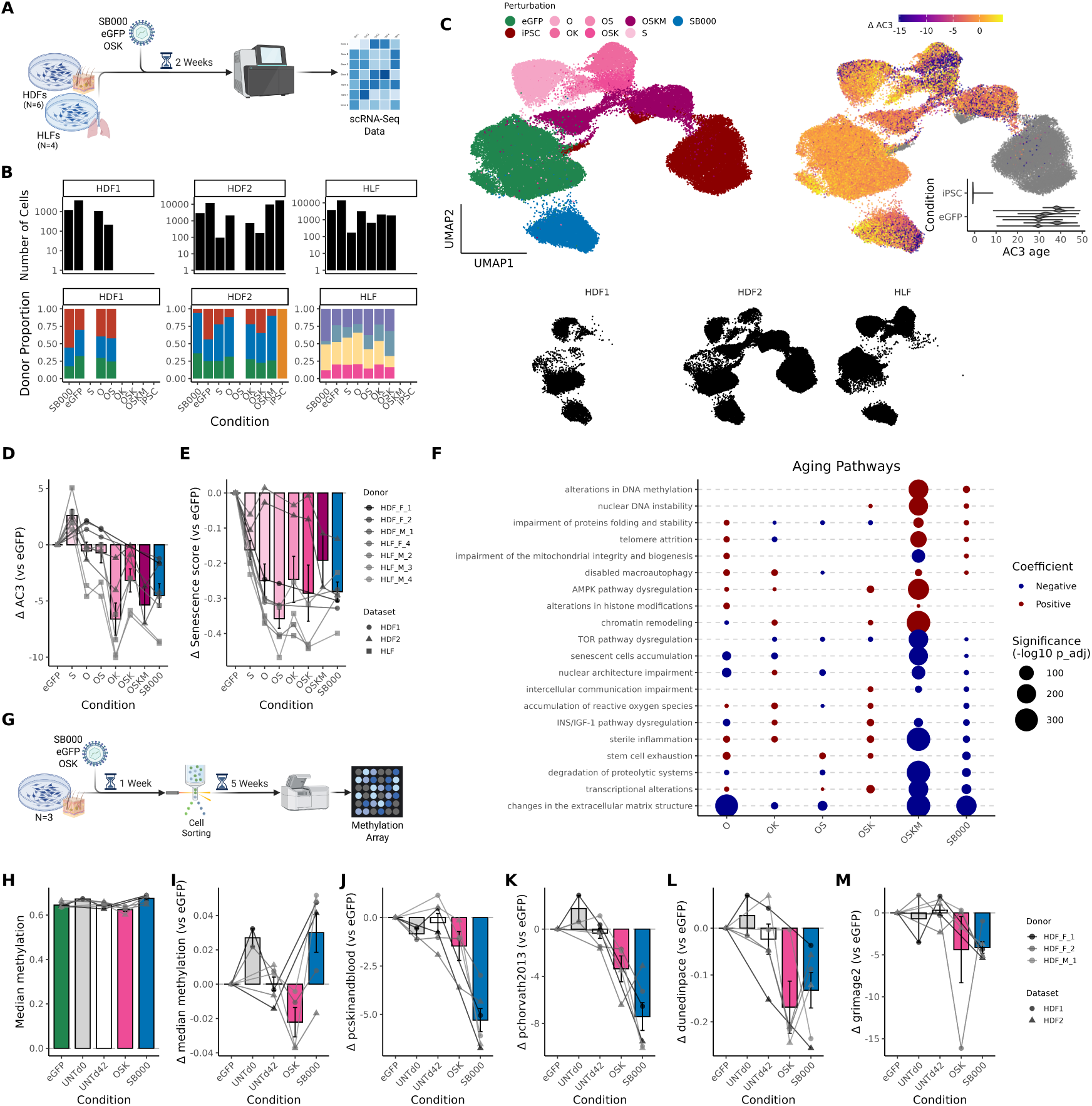
Multi-omic rejuvenation of aged human fibroblasts. **(A)** SB000 and the individual Yamanaka factors O, S and K were encoded in lentiviral vectors linking expression of the genes to eGFP. Human dermal fibroblasts (HDFs) and human lung fibroblasts (HLFs) from aged donors were exposed to lentiviruses expressing these vectors, eGFP-only vector, no virus or Sendai-virus encoded OSKM. Samples were analyzed by scRNA-seq two weeks after transduction, alongside an induced pluripotent stem cell (iPSC) control. **(B)** shows the contribution of donors and fibroblast organ from each experiment to the merged dataset. **(C)** The merged gene expression data were used to generate a UMAP projection, where cells are coloured by perturbation (left), or predicted transcriptomic age according to AC3 (right), with iPSCs as a positive control (predicted age *≈* 0, inset). The contribution of each experiment to the merged dataset is shown in silhouette (bottom). **(D)** Mean age prediction for rejuvenation interventions predicted by our transcriptomic clock, AC3, relative to eGFP control. **(E)** Mean expression of the SenMayo transcriptomic senescence score, relative to eGFP control. **(F)** Associations between perturbation probability and Open Genes ageing pathway scores in single cells (see methods). Pathways with a positive enrichment coefficient (i.e. gene expression is increased, consistent with a pro-aging effect) are colou6red red, whereas pathways with a negative enrichment coefficient (consistent with a reduced aging phenotype) are coloured blue. **(G)** In parallel, cells were sorted for eGFP positivity and maintained in culture. Six weeks post-transduction, genomic DNA was isolated from bulk samples and analysed for CpG methylation using the EPICv2 array. **(H)** shows the median proportion of methylated CpG sites throughout the genome in each sample, with **(I)** showing the changes in median methylation level relative to eGFP control. **(J-M)** The epigenetic age of cells expressing SB000, OSK or eGFP is shown according to PCSkinAndBlood **(J)**, PCHorvath2013 **(K)**, DunedinPACE **(L)**, or GrimAge2 **(M)** relative to eGFP control. In panels **D**, **E** and **H**-**M**, each point shows the mean value for cells from a single donor, in a single dataset, and bars and error bars show the mean and standard error for each condition. Lines link samples from the same experiment.

To investigate whether the observed decrease in transcriptomic age coincided with attenuation of cellular senescence, we calculated the SenMayo senescence signature [41]. This score exhibits only a weak correlation with chronological age across the AC3 training dataset (Pearson = 0.2, p = 0.03; data not shown), suggesting it provides a quasi-orthogonal measure of an age-related phenotype. SB000 significantly reduced the score (Figure 2E), achieving a mean reduction of 0.281 (s.d. 0.0780 p = 1.89 *×* 10*^−^*_5_), comparable to OSK’s 0.285 decline. OSKM caused a 0.158 decrease in the mean senescence score (0.0955 s.d., p = 0.103). Notably, the response to OSKM was heterogenous, with individual cells spanning a large range of negative and positive changes in senescence score, whereas SB000 produced a more homogeneous reduction (Figure S1). These findings demonstrate that SB000 achieves a consistent repression of the senescence programme across treated cells.

To explore the transcriptional pathways underlying SB000’s effects, we analysed the enrichment of the Open Genes ageing pathways in the expression profiles of perturbed cells [42]. SB000 significantly modulated 11 of 20 hallmarks in a beneficial direction in perturbed cells, while detrimentally shifting expression in 6 of 20 pathways, with the strongest positive effects observed in extracellular-matrix organisation, inflammatory signalling, proteolysis, and nuclear-architecture maintenance (Figure 2F). Although OSKM had a number of advantageous effects on transcriptomic ageing pathways (10 of 20), it simultaneously enriched for detrimental programmes (8 out of 20) such as DNA instability and chromatin remodelling. Despite lower significance levels compared to the full cocktail, omission of c-MYC increased the relative number of negative effects (2 of 20 beneficial, 9 of 20 detrimental), with removal of two or more factors displaying a variable impact. These results indicate that SB000 reproduces the beneficial aspects of OSKM while circumventing many of its apparent heterogenous pro-ageing effects.

To validate whether SB000-mediated rejuvenation extends beyond the transcriptome, we quantified global CpG methylation in SB000-treated HDFs from the same donors, as global CpG methylation is known to decline during ageing due to loss of heterochromatin [43, 44, 45, 46]. After six weeks of constitutive expression (Figure 2G), cells expressing SB000 exhibited a 3.00% higher median global methylation than eGFP controls (2.8% s.d., p = 0.0473) and similar levels to untreated samples that had been collected before the experiment, consistent with the maintenance of a more youthful epigenome (Figures 2H, 2I). OSK, on the other hand, tended to decrease global methylation (−2.21% mean, 1.71% s.d., p = 0.0817), likely due to the generalised chromatin opening seen during induction of pluripotency [47]. The CpG methylation level in binding sites for both AP-1 (previously implicated as a driver of age-associated chromatin opening [48]) and PRC2 (binding of which is thought to protect against ageing [49]) increased alongside the global hypermethylation with SB000 when compared to eGFP (Figures S3A, S3B). This suggests that SB000 does not act via differential methylation activity at the binding sites of either complex.

To further examine SB000’s effect on the epigenome, we computed epigenetic age according to DNAm ageing clocks in the same samples (Figure 2G). SB000 significantly reduced age estimates from four well-validated clocks, PCSkinAndBlood, PCHorvath2013, GrimAge2, and DunedinPACE, with PCHorvath2013 exhibiting a striking decline of 7.42 years (Figures 2J to 2M) [33, 50, 51]. The mean magnitude of these reductions matched or exceeded those achieved by OSK across all four clocks (PCSkinAndBlood −5.29 vs −1.47, PCHorvath2013 −7.42 vs −3.36, GrimAge2 −4.12 vs −4.38, DunedinPACE −0.132 vs −0.168). SB000 actively reversed epigenetic ageing rather than slowed its progression in culture as shown by the similar reduction in age compared to untreated samples at day zero [52]. Quantitative analysis of an expanded panel of over 20 clocks, including causal and stochastic clocks, revealed consistent rejuvenative trends (Figure S2). The epigenetic age reductions were not driven by global hypermethylation, as indiscriminately increasing methylation values artificially produced mixed changes in epigenetic clocks (data not shown). Collectively, these data establish SB000 as a potent modifier of epigenetic age.

### SB000 is safer than the Yamanaka factors

The Yamanaka factors were identified by explicitly maximising pluripotency, an objective that inevitably erodes specialised cellular functions and carries substantial teratogenic risk [21]. We therefore hypothesised that a rejuvenation gene discovered and optimised for biological age reversal rather than de-differentiation could better preserve fibroblast identity while avoiding pluripotency. To test this proposition, we bench-marked SB000 against the Yamanaka factors (and combinations thereof) across transcriptomic, epigenomic, and functional assays designed to expose threats to cellular identity.

In order to determine whether SB000 perturbs fibroblast lineage commitment, we first quantified a simple transcriptomic signature of fibroblast identity [53]. While SB000-treated cells exhibited a small reduction in fibrobast identity score compared to eGFP controls (mean score change −0.594, s.d. 0.316, p = 0.00111), the expression of any Yamanaka subset was associated with larger reductions, the greatest of which was observed after OSKM expression (Figures 3A, S5). This included a significant reduction for OSK-expressing cells (mean score change −2.13, s.d. 0.235, p = 3.48*×*10*^−^*_6_). Analysis of the five individual identity genes revealed patterns of expression that were largely similar between SB000- and eGFP-treated cells, for example in the strong expression of genes such as *DCN* and *LUM*. The subtle reduction in mean fibroblast score with SB000 was driven by reduced expression of the single gene *PDGFRA* (Figure 3B). These data indicate that SB000 leads to less disruption in the core transcriptional programme of fibroblasts than the Yamanaka factors.

**Figure 3:**
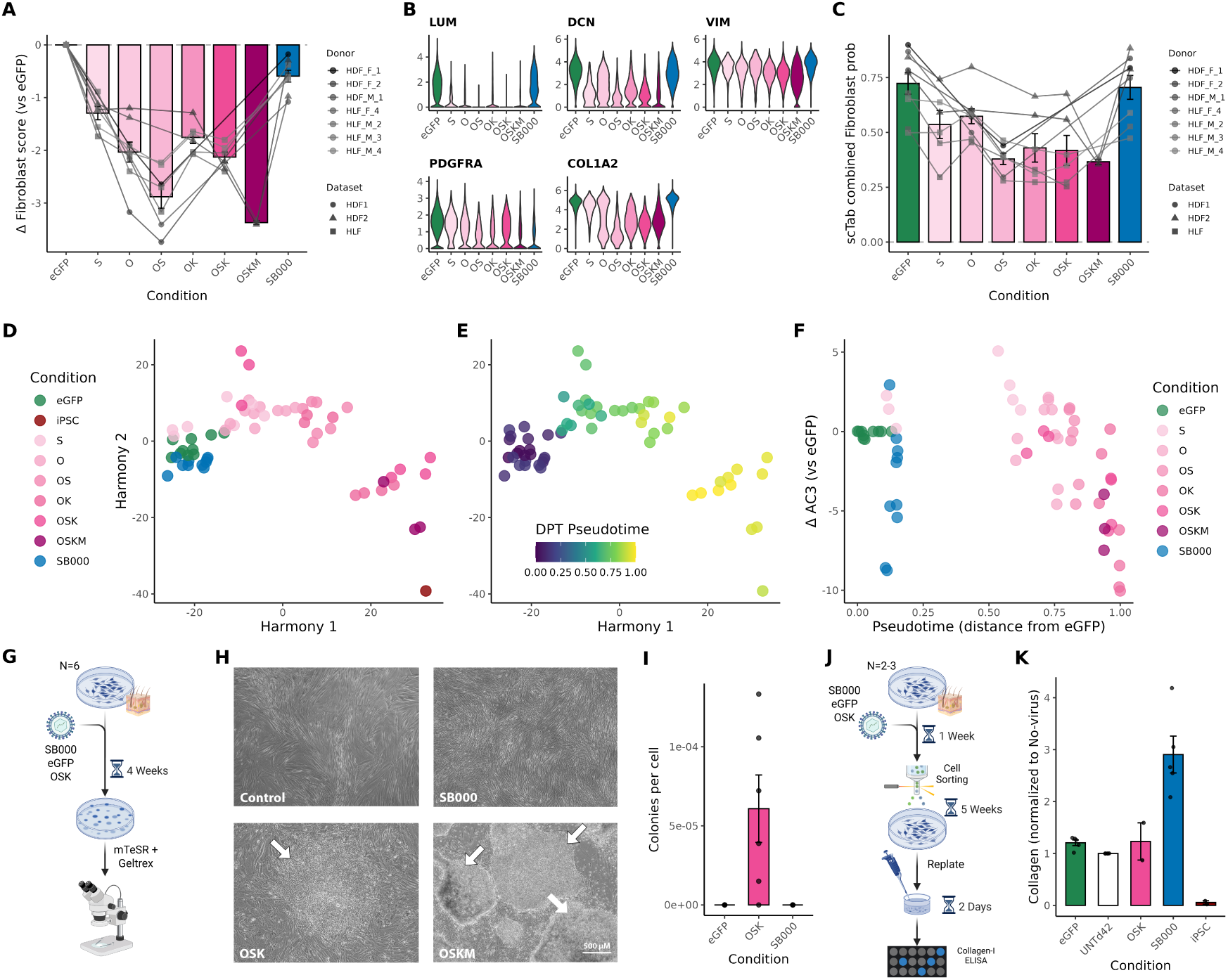
Maintenance of cellular identity in SB000-rejuvenated cells. Human dermal fibroblasts (HDFs) expressing SB000, subsets of the Yamanaka factors or eGFP control were analysed by scRNA-seq two weeks post-transduction. The expression of known fibroblast identity genes was used to calculate an identity score of single cells. **(A)** The mean fibroblast score of each sample, relative to eGFP control. **(B)** Expression of the five constituent genes from the score in (A). **(C)** The mean probability of cells being classified as fibroblasts by the cell type classifier model scTab. **(D-F)** The pseudobulked single-cell transcriptomic data were used to calculate a pseudotime trajectory. Samples are coloured by the intervention they received **(D)** or their progression in pseudotime **(E)**, and their relative transcriptomic age compared to the eGFP control is plotted vs. pseudotime **(F)**. **(G)** HDFs from the same aged donors were then transduced with the lentiviral vectors and maintained in culture on GelTrex (without sorting), and the morphology monitored. **(H)** Representative images of the colony formation assay after four weeks. Colonies are shown with white arrows. **(I)** The number of colonies was enumerated for each sample, and the number normalised to the number of cells that were eGFP positive. **(J)** Cells were replated at a fixed density six weeks after transduction and their medium assayed for Collagen I. **(K)** shows the concentration of collagen I in the supernatant of cells, after normalisation to the total protein content, and expressed relative to the concentration in samples from cells without any virus. In panels **A & C**, each point shows the mean value for cells from a single donor, in a single dataset, and bars and error bars show the mean and standard error for each condition. Lines link samples from the same experiment.

To corroborate the maintenance of fibroblast gene expression, we next applied the highly accurate single-cell annotation model scTab [54]. SB000 did not significantly change the proportion of cells confidently labelled as fibroblast-like, maintaining a probability close to that seen in eGFP controls (mean 70.5% vs. 75.0%, p = 0.151), whereas the Yamanaka cocktails reduced classification confidence, roughly in proportion to the number of factors present (OSK mean 41.8%, p = 0.00265; Figure 3C). The alignment between gene-signature and scTab metrics strengthens the evidence that SB000 does not compromise transcriptomic markers of somatic identity.

To provide an unsupervised comparison of the interventions, we positioned SB000-, Yamanaka-, and eGFP-treated scRNA-seq samples within a batch-corrected PCA derived from iPSC and fibroblast reference datasets. Pseudobulk projections revealed overlap of SB000 samples with eGFP controls, whereas progressive inclusion of Yamanaka factors shifted transcriptomes along a de-differentiation trajectory toward iPSCs, with OSKM exerting the strongest effect (Figure S6). A quantitative pseudotime score mirrored this gradient, placing SB000 and eGFP at the somatic terminus and all Yamanaka cocktails at increasingly advanced positions along an apparent pluripotency and de-differentiation axis (Figures 3D, 3E). There was no correlation between pseudotime and AC3-predicted transcriptomic age in this analysis: remarkably, SB000 treatment drove equivalent transcriptomic rejuvenation to OSKM while treated cells remained anchored to their fibroblast identity (Figure 3F). This analysis suggests that SB000 has decoupled robust transcriptomic rejuvenation from pluripotency.

To provide functional confirmation that SB000 does not induce pluripotency, we performed a colony-formation assay that reports reprogramming efficiency (Figure 3G). Samples transduced with OSKM formed pluripotent colonies as expected. Colonies also formed in samples transduced with lentiviruses encoding O, S and K individually, despite the omission of c-MYC, as reported previously [31]. By contrast, SB000 never produced colonies across multiple independent replicates, and the monolayers retained a fibroblast-like morphology throughout extended culture (Figures 3H, 3I). These results provide phenotypic confirmation that SB000 does not trigger a reprogramming cascade.

Finally, seeking a fibroblast-specific functional read-out, we measured secretion of collagen I, the production of which declines with age (Figure 3J) [55, 56]. Cells expressing SB000 secreted significantly higher levels of collagen I than eGFP controls, whereas iPSC cultures derived from HDFs produced negligible amounts (Figure 3K). OSK-positive cells secreted collagen at control-like levels, confirming partial identity retention but without any enhancement (Figure 3K). These observations reveal that SB000 safeguards a canonical fibroblast function.

### SB000’s rejuvenation generalises across germ layers

Much of the promise of the Yamanaka factors arises from their ability to rejuvenate disparate cell types. We therefore investigated whether SB000-mediated rejuvenation is similarly generalisable. To span two of the three germ layers, we gathered ectoderm-derived normal human epidermal keratinocytes (NHEKs) to supplement the existing data from mesoderm-derived fibroblasts. We computed epigenetic age six weeks after transduction with SB000, eGFP, or OSK in NHEKs from two aged female donors (Figure 4A).

**Figure 4:**
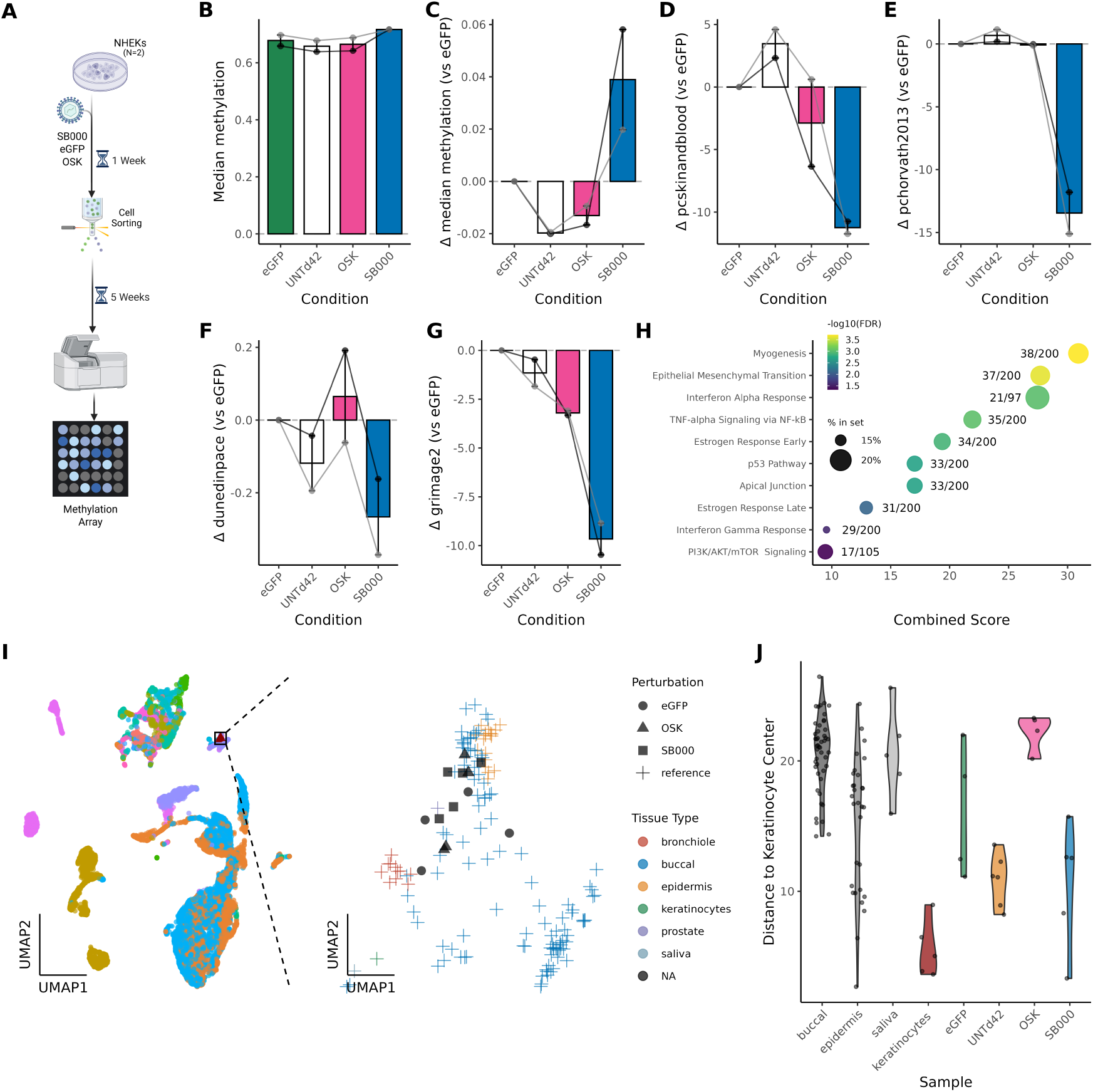
Multi-germ layer epigenetic rejuvenation by SB000. **(A)** Normal human epidermal keratinocytes (NHEKs) were transduced with SB000, OSK or eGFP control lentivirus, and the fluorescence population sorted using FACS. Six weeks post-transduction, the epigenome of bulk samples was profiled using the EPICv2 array. **(B)** shows the median methylation levels across the array CpG sites in SB000-treated NHEKs, and **(C)** the change in median methylation level, relative to eGFP control. **(D-G**) The bulk epigenetic age was calculated according to **(D)** PCSkinAndBlood, **(E)** PCHorvath2013, **(F)** DunedinPACE or **(G)** GrimAge2 (shown relative to eGFP control). **(H)** shows the pathways in which differentially hypermethylated regions were enriched in SB000-treated NHEKs, relative to eGFP control. **(I)** the DNAm profiles of NHEK samples expressing the rejuvenation genes were mapped onto a large, multi-tissue reference dataset (AltumAge) using CpGPT. The UMAP plot of the embedded dataset is shown, with perturbed samples highlighted. Inset: the local neighbourhood of the samples. **(J)** To refine tissue/cell-type similarities to each of our perturbations, we first performed hierarchical clustering on the top 20 PCs of the AltumAge dataset combined with our samples. Then, we performed PCA using the subset of samples that were within the same cluster that our data was part of. Finally, we me1a0sured distance using PC2 and PC3 (as PC1 captured a batch effect) to keratinocytes from the AltumAge data in the same cluster. In panels **B**-**G**, each point shows the mean value for cells from a single donor, in a single dataset, and bars and error bars show the mean and standard error for each condition. Lines link samples from the same experiment.

Assessing the global methylation landscape, we found that SB000 elicited an approximately 4% median increase in CpG methylation relative to eGFP controls at the six week timepoint, consistent with a more youthful epigenome (Figures 4B, 4C). OSK decreased global methylation compared with eGFP (approx. −1.5%), although levels were comparable to untreated cultures (Figure 4C). Consistent with these results, SB000-treated cells had higher levels of DNAm at both AP-1 and PRC2 binding sites than control cells, as seen in HDFs (Figures S8A, S8B). These observations suggest that SB000 maintains heterochromatin integrity in both ectoderm- and mesoderm-derived lineages.

To quantify epigenetic age we again applied the same four well-validated clocks — PCSkinAndBlood, PCHorvath2013, DunedinPACE and GrimAge2 — with PCHorvath2013 exhibiting a remarkable median decline of 13.6 years with SB000 (Figures 4D to 4G). The magnitude of the SB000-mediated age decrease in NHEKs exceeded that observed in HDFs, especially relative to OSK. Strikingly, the rate of change in PCHorvath2013 approached a decade of reversal per month of treatment (−9.7 years per month). Similar improvements were captured by dozens of additional clocks that predict chronological age, phenotypic age, mortality, mammalian age, intrinsic age, and causal epigenetic age (Figure S7). This cements SB000 as an exceptionally powerful, multi cell-type epigenetic rejuvenator.

To explore gene-specific epigenetic modulation we examined differentially methylated CpG sites with SB000 when compared to the eGFP controls. No clear enrichment in MSigDB Hallmark 2020 [57] pathways was observed among genes with hypomethylated promoters (i.e. potential expression increases). Several pathways, however, showed significant promoter hypermethylation, suggesting inhibition of gene expression (Figure 4H). Most notable were myogenesis, epithelial-to-mesenchymal transition (EMT), interferon-*α* response, and TNF-*α* signalling via NF-*κ*B. As inflammatory pathways are tightly linked to age-associated decline (inflammageing), the observed suppression of these programmes aligns with broad rejuvenation.

Analogous to the prediction of biological age based on epigenetic clocks, we aimed to predict the concentration of plasma proteins based on the methylome, as they are more easily interpretable disease biomarkers. CpGPT [58] was fine-tuned to estimate circulating levels of hundreds of proteins, 27 of which overlap a list of morbidity- and mortality-associated factors based on a recent UK Biobank survey [59]. While DNAm biomarker proxy models are not perfect predictors, they temporally integrate highly variable plasma proteome signals into a consistent metric [60, 61, 50]. We then applied this model on our data, revealing that SB000 treatment in NHEKs was associated with a decrease in the predicted levels of several such disease-associated factors, including CHI3L1, IL-6, and multiple TNF-associated proteins, indicative of reduced organismal inflammation (Figure S9A). When this analysis was repeated for the DNAm samples from fibroblasts described earlier (Figure S4), there was a significant and strong positive correlation between the predicted change in plasma proteins in the two cell types (SB000 Pearson = 0.847, OSK Pearson = 0.609; Figure S9B). Conservatively, this suggests that SB000-treated aged cells better resemble those that come from a body with lower disease and mortality risk. The predictions may also provide preliminary evidence that the benefits of SB000 would extend beyond directly treated cells.

Finally, to probe the effect of SB000 on keratinocyte identity, we projected whole-genome methylation profiles from the SB000-, OSK-, and eGFP treated samples into a biologically meaningful embedding space using the 100-million-parameter CpGPT foundation model [58]. When projected into the large DNAm dataset used to train AltumAge [62], the treated cells all clustered closely to samples from skin, suggesting that there had been no drastic changes in cell identity for either SB000- or OSK-expressing cells (Figure 4I). After correcting for batch effect, the distance from keratinocyte samples in the reference dataset to untreated cells and those treated with SB000 was similar in PCA space (Figures 4J, S10). Although less comprehensive than our transcriptomic and functional assays in fibroblasts, these data nevertheless suggest that SB000 preserves cell identity based on DNAm patterns while exerting potent, cross-germ-layer rejuvenation.

## Discussion

Rejuvenating human cells while preserving their specialised identity remains a central challenge in translational geroscience. To overcome this challenge, we developed one of the few machine learning platforms optimised specifically for the discovery of safe rejuvenation genes. Here, we leverage AC3, a highly accurate single-cell transcriptomic clock, to identify SB000, a single gene which rejuvenates the transcriptome of fibroblasts more than previously reported single genes [38, 39, 40] and rivals the potency of the Yamanaka factors without the liability of pluripotency. Two weeks of SB000 expression are sufficient to reverse transcriptomic age, reduce senescence signatures, and attenuate multiple hallmarks of ageing, thereby establishing a robust foundation for multi-omic rejuvenation. Six weeks of overexpression powerfully restores DNA methylation patterns to a youthful state.

To contextualise SB000’s performance, we compared its effects to those of OSKM, the gold-standard epigenetic rejuvenator. Starting with cellular senescence, quantitative single-cell analysis revealed that OSKM induces a highly heterogeneous senescence response, consistent with the fate bifurcation in which a minority of cells fully reprogram while the remainder enters proliferative arrest [63]. By contrast, SB000 produces a homogeneous and significant reduction in the SenMayo signature, indicating that targeted modulation of a single gene can elicit uniform anti-senescent effects without triggering divergent cell fates.

Functional rejuvenation of fibroblasts was supported by collagen I secretion assays. Age-related decline in collagen production is well documented, with dermal fibroblasts from elderly donors synthesising significantly less collagen I than their young counterparts [55, 56]. Whereas iPSCs cultures secreted negligible collagen, SB000 expression maintained collagen output and even enhanced it. The specific clinical implications of the increase in collagen secretion remains to be explored. Regardless, these data strongly support that SB000 preserves a key fibroblast-specific function, providing a phenotypic correlate to the molecular rejuvenation captured by the transcriptomic and epigenetic clocks.

Achieving genuine epigenetic rejuvenation in primary human cells is notoriously difficult, especially within genetically heterogeneous populations. Numerous studies claiming multi-omic age reversal did not report the effect on gold-standard epigenetic clocks [38, 64, 39, 40]. SB000 overcomes this barrier by reducing age estimates across dozens of well-validated clocks, including PCSkinAndBlood, PCHorvath2013, GrimAge2, and DunedinPACE. In keratinocytes, SB000 decreased the PCHorvath2013 metric by close to ten years for each month of treatment and lowered the DunedinPACE pace-of-ageing score by approximately 25%. This establishes SB000 as, to our knowledge, the most powerful epigenetic age-reversal gene reported to date for human cells. Crucially, longitudinal sampling demonstrated that SB000 drives absolute age reversal rather than merely slowing culture-induced epigenetic drift.

While the mechanism of SB000-driven DNA methylation is yet to be explored, the observed genome-wide hypermethylation inclusive of high PRC2-binding sites is intriguing. It has been hypothesized that rejuvenation interventions act at least partially through increasing accessibility at bivalent chromatin regions [49]. OSKM has been shown to decrease methylation at these loci, making them similar to embryonic cells. Here, we observe broad epigenetic rejuvenation without hypomethylation of PRC2-binding sites, indicating that loss of cell identity and acquisition of stem-like characteristics may not be necessary for reversal of the ageing phenotype.

Safety considerations further elevate the therapeutic potential of SB000 above current state-of-the-art interventions. Unlike OSKM, which requires pulsed or transient dosing to minimise teratogenic risk, SB000 maintains normal expression of fibroblast identity markers such as *DCN* and *LUM* and does not drive colony formation in reprogramming assays. The single-gene format also streamlines the potential path to a therapeutic by eliminating stoichiometric constraints during vector manufacture, simplifying dose optimisation and facilitating mechanistic dissection of the rejuvenation pathway.

Evidence that SB000 acts through a core anti-ageing mechanism is provided by its cross-tissue efficacy. The gene reverses molecular age in mesoderm-derived dermal and lung fibroblasts as well as ectoderm-derived keratinocytes, implying that its downstream network operates across germ-layer boundaries.

SB000 was discovered by optimising directly for age reversal, rather than for pluripotency, using a machine-learning-guided, single-cell-resolution screening platform. SB000 charts a tractable route toward safe, durable, and broadly applicable rejuvenation. Future work with this intervention will delineate the molecular circuitry that orchestrates its rejuvenative effects, laying the groundwork for next-generation therapeutics aimed at extending human healthspan and lifespan.

## Limitations

### Functional rejuvenation

The transcriptome and epigenome provide compelling evidence that SB000 rejuvenates human cells independent of germ layer, through ageing clock reversal and improvement in gene expression signatures linked to the hallmarks of ageing. SB000 also increases secretion of collagen I, suggesting that it may rescue age-related functional decline. However, further work is required to determine the scope of functional rejuvenation in fibroblasts and beyond.

### Safety

Transcriptomic analysis, collagen secretion and a colony formation assay suggest that SB000-rejuvenated fibroblasts maintain identity without progressing to pluripotency. However, the safety of a therapeutic is also governed by factors beyond cell identity and pluripotency, so progression to the clinic requires ongoing and robust safety monitoring *in vivo*.

## Methods

### Human dermal fibroblast transductions

HDFs were cultured in 2D using standard media conditions. For arrayed single-cell RNA-seq screens, lentiviral vectors (LVs) carrying the genes of interest (GOIs) were first arrayed individually into 96-well plates and stored until the experiment day. Human dermal fibroblasts were plated into 96-well plates and transduced 24 hours later by transferring the LV array plate contents into their corresponding screening plates. The transduced cells were maintained with media changes twice per week. On day 15, each screening plate was harvested by pooling all wells, resulting in 19 pools of 80 GOIs spiked with an eGFP control. Each pool was then fixed for single-cell RNA sequencing (scRNA-seq). For DNAm clock confirmation, HDFs were plated into 6-well plates and transduced with LVs carrying eGFP-tagged GOIs after 24 hours. On day 7, transduced cells were harvested and the eGFP+ population was isolated by FACS and maintained with full media changes three times per week. On day 42, HDF cells were harvested and processed to extract genomic DNA.

### Single-cell RNA-seq

Single-cell RNA-seq library preparation was performed using the ScaleBio Single Cell RNA Sequencing kit as per the manufacturer’s instructions.

### Transcriptomic data analysis

Raw single-cell reads were aligned both to the human genome hg38 using ENSEMBL’s 112 gene annotation and to our exogenous vectors with STARSolo [65]. Generated count matrices were filtered against cells with less than 500 unique genes, 1500 counts, and over 20% of mitochondrial DNA reads.

Fibroblast scores were calculated with `scanpy’s scanpy.tl.score_genes` function with different subsets of genes. It was derived from a subset of five genes (LUM, DCN, VIM, PDGFRA, COL1A2) whose expression is able to differentiate fibroblast from other skin cell types [53]. The senescence score is calculated from the SenMayo gene set [41].

Cell type prediction was done with scTab using the default parameters [54]. All cell type labels with “fibroblast” in the name were grouped together.

### Measuring DNA methylation

Genomic DNA was isolated from cell pellets using the DNeasy Blood & Tissue kit (Qiagen #69506). gDNA was submitted for DNA methylation analysis with the Illumina EPICv2 array.

### DNA methylation data analysis

Processed beta matrices were obtained with the R package `sesame`. `openSesame` was used with the default parameters. Samples with less than 70% of *frac_dt*, the percentage of detected probes, were removed due to failed quality control. Technical replicates per donor were averaged together for analysis.

All epigenetic clocks were calculated with `pyaging` [66]. For the computation of the mammalian clocks, the beta values in the mammalian array were imputed using the 100-million parameter CpGPT. Otherwise, KNN imputation was used.

Mortality-associated DNA methylation protein proxies were calculated with CpGPT-2M-Proteins [58]. The proteins were chosen by the intersection of the 322 plasma proteins predicted with the foundation model and 54 mortality- and disease-associated proteins from proteomic analysis of the UK Biobank [59], yielding a total of 27 proteins. Predictions of the 322 proteins were mean-normalised per sample.

Our methylation samples were mapped using the 100-million parameter CpGPT to the AltumAge dataset with the faiss package [58, 62, 67].

Differentially methylated CpGs were calculated for SB000 against eGFP using a generalised linear model with donor as a covariate. CpG sites with missing values were filtered. P-values were Bonferroni-corrected with the number of remaining CpG sites across all samples. Sites with a p-adjusted value above 0.05 that fell within 1500 basepairs of the transcription start site were assigned to a gene. Genes were further divided into two groups depending on whether the change in methylation was positive (hypermethylation) or negative (hypomethylation). Pathway enrichment for the MSigDB Hallmark 2020 database [57] for the two lists was calculated with `gseapy`.

DNA methylation enrichment in binding sites was done by averaging the log-fold change in signal of the ENCODE ChIP-Seq samples with the AP-1 subunits JUN (ENCFF661FPB), FOSL1 (ENCFF753AIX), JUND (ENCFF940XSY), ATF2 (ENCFF031CBV), and ATF3 (ENCFF753CFJ). Beta values for each perturbation were aggregated and ranked based on the average AP-1 signal. A similar analysis was done for the PRC2 complex based on the EZH2 (ENCFF105JFX) and SUZ12 (ENCFF224RZW) subunits, as per [49].

### Supernatant collagen assay

On day zero, LV-transduced HDFs and controls were harvested and seeded into 6-well plates. Plates were incubated at 37*^◦^*C for 48 hours. The conditioned media was then collected, clarified by centrifugation (2500 rpm at 4*^◦^*C for 5 min), and stored at −80*^◦^*C. In parallel, cell pellets were harvested and stored at −80*^◦^*C until the day of the assay. On the day of the assay, clarified media and cell pellets were thawed on ice. Pellets were lysed in RIPA Buffer (Thermo #89900) following the manufacturer’s protocol. Total protein concentration from media and lysates was quantified using a BCA Protein Assay (Thermo #23225), and type-I Collagen levels were quantified using the Human Collagen Type-I ELISA Kit (Abcam #ab285250). Absorbance was measured using a HidexSense plate reader.

### Pluripotent colony formation assay

HDFs were seeded at 200,000 cells per well of a tissue culture 6-well plate (Corning, 10110151). On the following day, HDFs were transduced with individual lentiviruses encoding the SB000 or single Yamanaka factors. Lentiviral transduction was performed by adding lentivirus to fibroblast media supplemented with polybrene (Millipore, TR-1003-G) at 8 *µ*g/mL, followed by spinfection in a centrifuge at 800 *g* for 30 min at 32*^◦^*C. Additional controls included a polybrene-only negative transduction control, as well as CytoTune-iPS 2.0 Sendai OSKM Reprogramming control (Invitrogen, A16517). Sendai transduction was performed without spinfection or polybrene, as described in the manufacturer protocol for Lot L2200077. 24 hours post-transduction, media was replaced with fresh fibroblast media. From then on, cell culture media was changed every other day. On day 5, a set of fresh 6-well plates was pre-coated with geltrex (Gibco, A14133-02), diluted at 1:100 in ice-cold DPBS (Life Technologies, 14190-144), followed by incubation overnight at 4*^◦^*C. On day 6, geltrex solution was aspirated from the coated plates. Then, each condition was passaged 1:1 into geltrex-coated plates in fibroblast media. On day 7, media was changed to mTeSR-Plus (Stem-Cell Technologies, 100-0274) supplemented with FGF2-containing mTeSR-Plus 5X Supplement (StemCell Technologies, 100-0275). From then on, all conditions were grown in complete mTeSR-Plus media; media changes were performed every other day. On day 26, all conditions were imaged and iPSC colonies were counted.

### Normal Human Epidermal Keratinocyte transduction

NHEKs were seeded into collagen-coated 6-well plates and left to recover for 1 week. Cells were then transduced using spinfection and incubated for 24 hours. Media was changed every other day for seven days, at which point transduced cells were selected using FACS. After sorting, cells were seeded in collagen-coated 6-well plates, and media changed every other day. Cells were not passaged between sorting and collection at day 42.

## Code and data availability

Data, code and materials will be made available upon publication.

## Acknowledgments

We are grateful to the UCL Genomics facility for collecting the EPICv2 array data in this work, and to the flow cytometry facility in the School of Biological Sciences, University of Cambridge. We also thank our scientific advisory board for their thoughtful feedback on the manuscript. Schematic diagrams were generated using stock images from Biorender.

## Supplementary Figures

**Figure S1:**
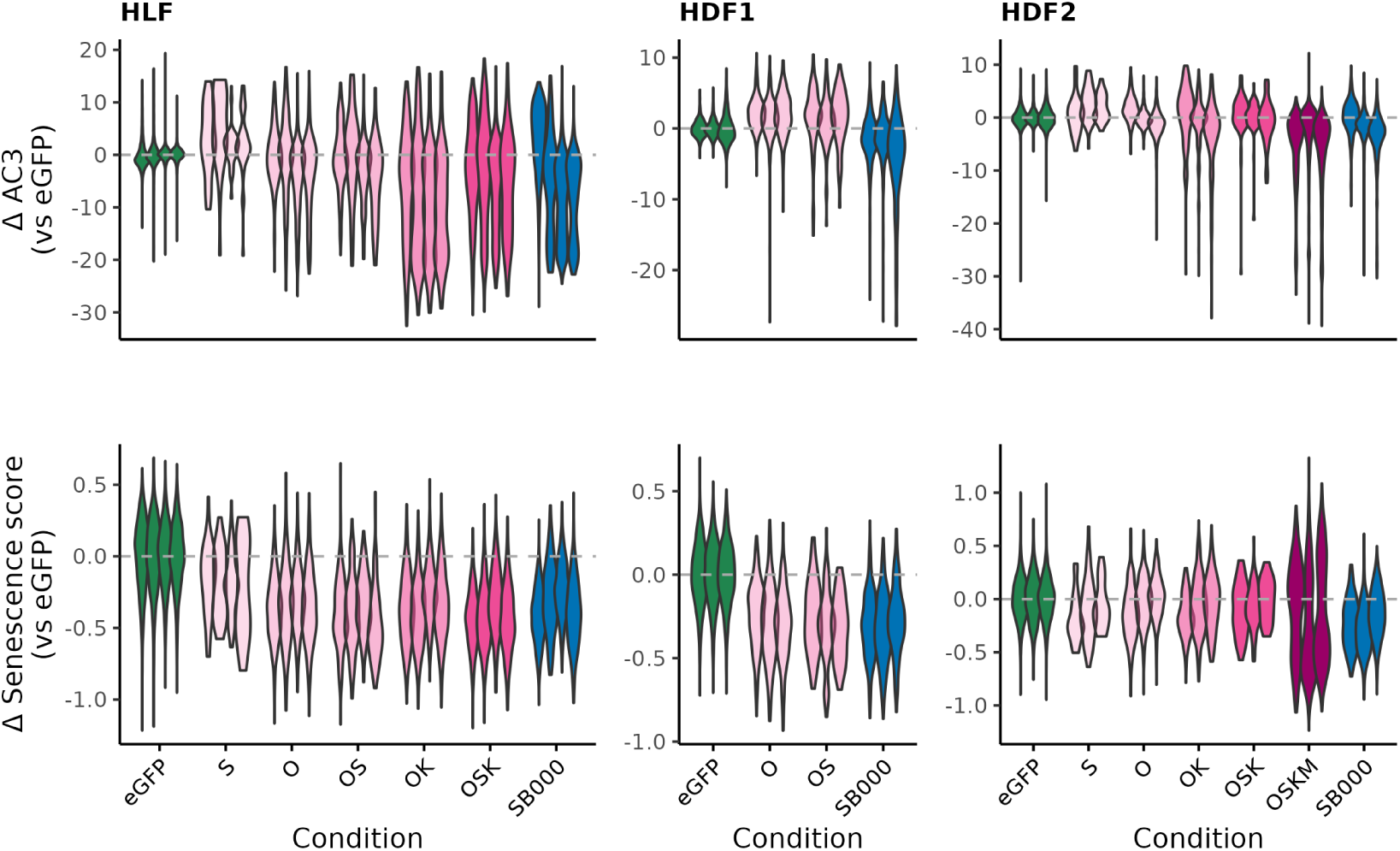
SB000 and the individual Yamanaka factors O, S and K were encoded in lentiviral vectors linking expression of the genes to eGFP. Human dermal fibroblasts (HDFs) and human lung fibroblasts (HLFs), both from aged donors, were exposed to lentiviruses expressing these vectors, eGFP-only vector, no virus or sendai-virus encoded OSKM. Top: the distribution of transcriptomic age in the single cells two weeks after receiving the different combinations of lentiviruses, according to AC3. Bottom: the distribution of SenMayo senescence scores for the same samples. Each column represents an independent experiment, one performed in HLFs and the other two in HDFs (“HDF1” and “HDF2”). Violins are coloured by condition, with separate violins for cell populations from different donors. Values are expressed relative to the median of eGFP samples from the same donor.

**Figure S2:**
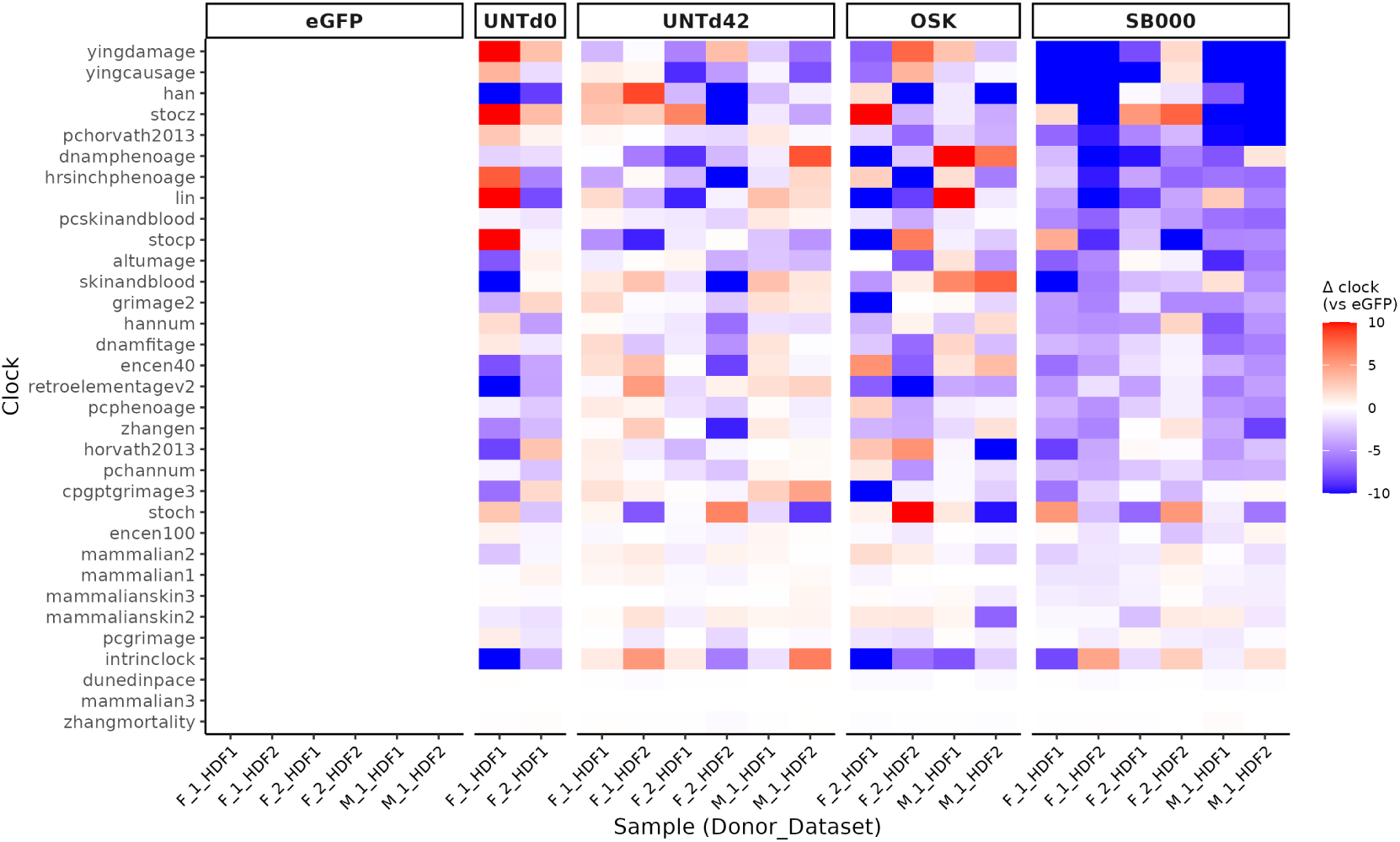
HDFs expressing rejuvenation genes were maintained in culture for six weeks, after which the DNAm status of their genomic DNA was profiled by EPICv2 array. The bulk epigenetic age of samples was calculated using the *pyaging* package; shown is the change in predicted age of cell samples receiving SB000, OSK, relative to eGFP-treated cells from the same donor, according to many published epigenetic clock algorithms. The clock results for untreated cells before the start of the experiment (“UNTd0”) and from the six-week time point (“UNTd42”) are included for reference.

**Figure S3:**
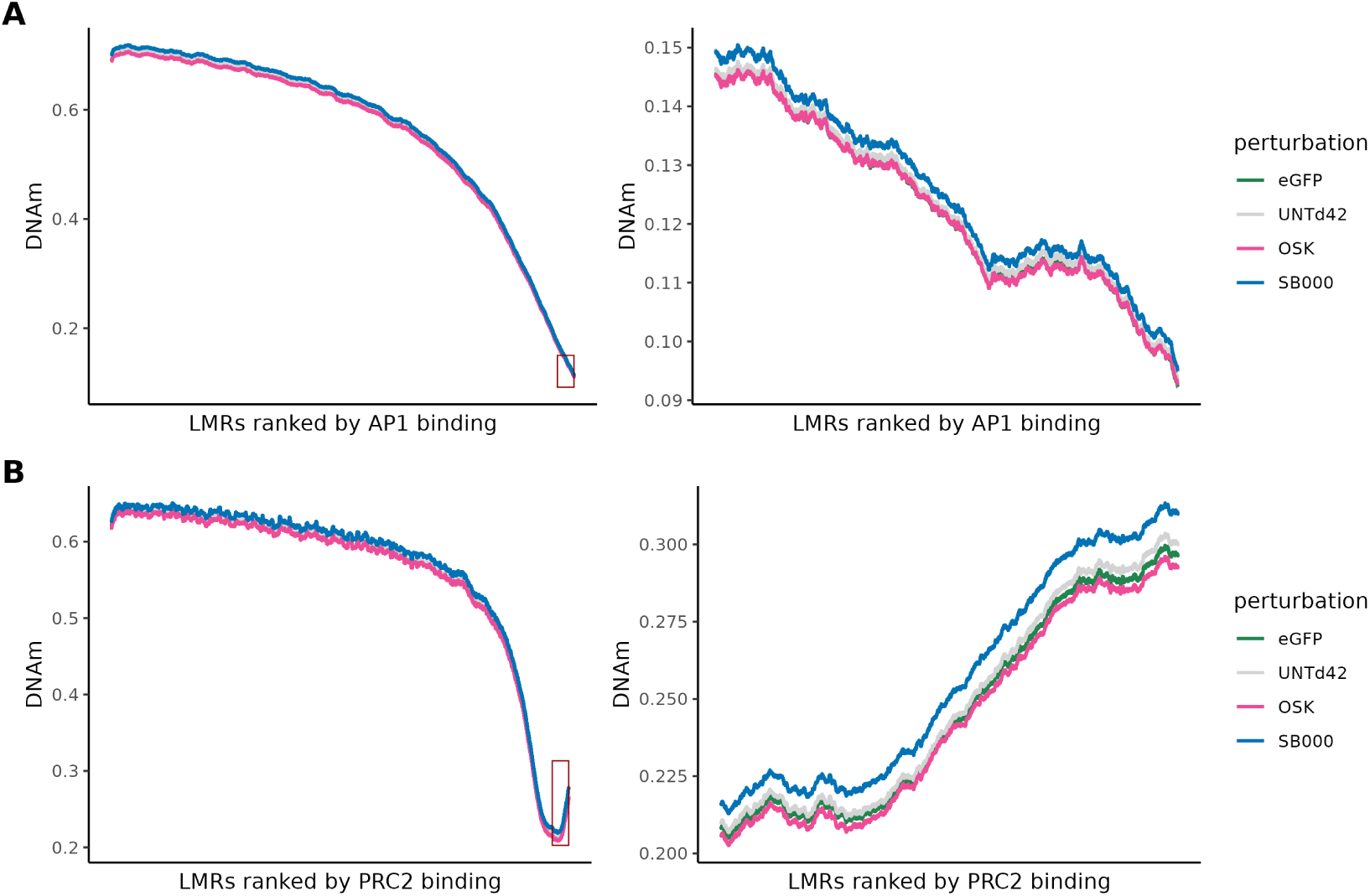
HDFs expressing rejuvenation genes were maintained in culture for six weeks, after which the DNAm status of their genomic DNA was profiled by EPICv2 array. **(A)**, left: the methylation status of DNA in AP1 binding sites (low-methylated regions, LMRs) across the methylome. The DNAm level at CpG sites is plotted in ascending order of AP1 binding enrichment, with each of the conditions coloured. Right: a zoomed representation of the top 1000 sites most enriched for AP1 binding (highlighted by a red box in the left panel). **(B)**, left: the methylation status of DNA in PRC2 binding sites (LMRs) across the methylome. The DNAm level at CpG sites is plotted in ascending order of PRC2 binding enrichment, with each of the conditions coloured. Right: a zoomed representation of the top 1000 sites most enriched for PRC2 binding (highlighted by a red box in the left panel).

**Figure S4:**
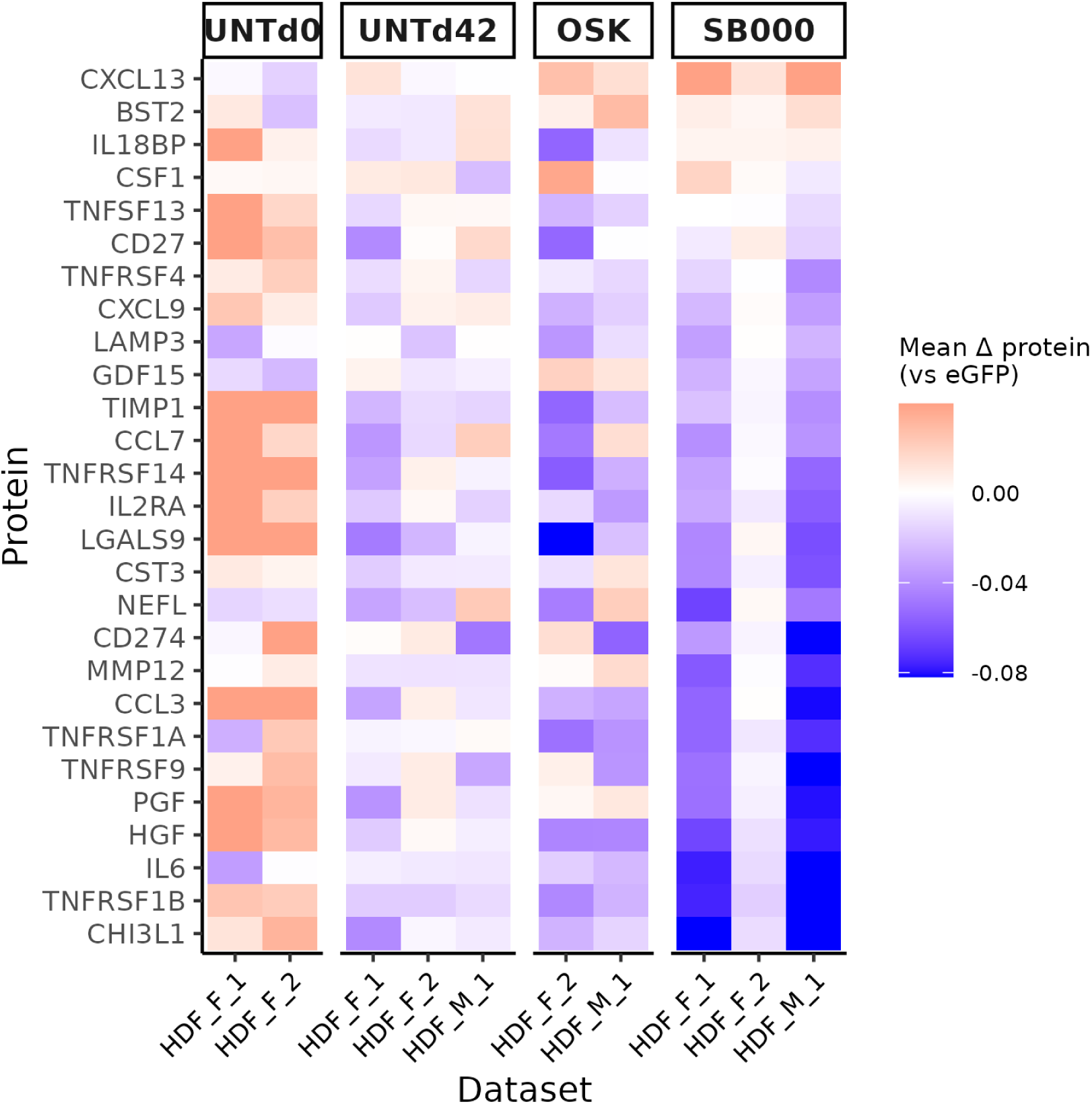
SB000 and the individual Yamanaka factors O, S and K were encoded in lentiviral vectors linking expression of the genes to eGFP. Human dermal fibroblasts (HDFs) and human lung fibroblasts (HLFs), both from aged donors, were exposed to lentiviruses expressing these vectors, GFP-only vector, no virus or sendai-virus encoded OSKM and the fluorescence population sorted using FACS. After six weeks, CpGPT was used to predict the level of circulating mortality-associated proteins from the DNAm profiles of these samples. The heatmap shows the predicted levels of disease-associated blood proteins for untreated cells before the start of the experiment (“UNTd0”) and from the six-week time point (“UNTd42”), as well as those expressing OSK or SB000. Values are expressed relative to those for eGFP samples from the same donor.

**Figure S5:**
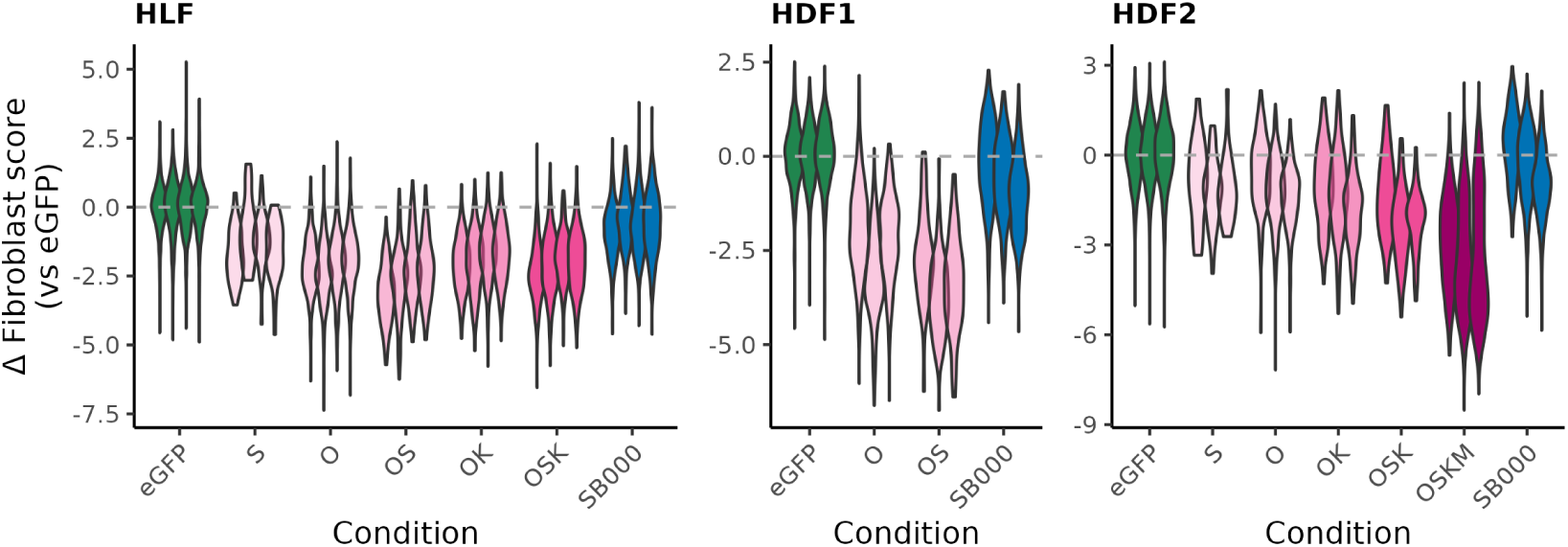
SB000 and the individual Yamanaka factors O, S and K were encoded in lentiviral vectors linking expression of the genes to eGFP. Human dermal fibroblasts (HDFs) and human lung fibroblasts (HLFs), both from aged donors, were exposed to lentiviruses expressing these vectors, eGFP-only vector, no virus or sendai-virus encoded OSKM. The violin plots show the distribution of fibroblast identity scores across single cells in the populations receiving the intervention, after two weeks. Each column represents an independent dataset, one in HLFs and the other two in HDFs (“HDF1” and “HDF2”). Violins are coloured by condition, with separate violins for cell populations from different donors. Values are expressed relative to the median of eGFP samples from the same donor.

**Figure S6:**
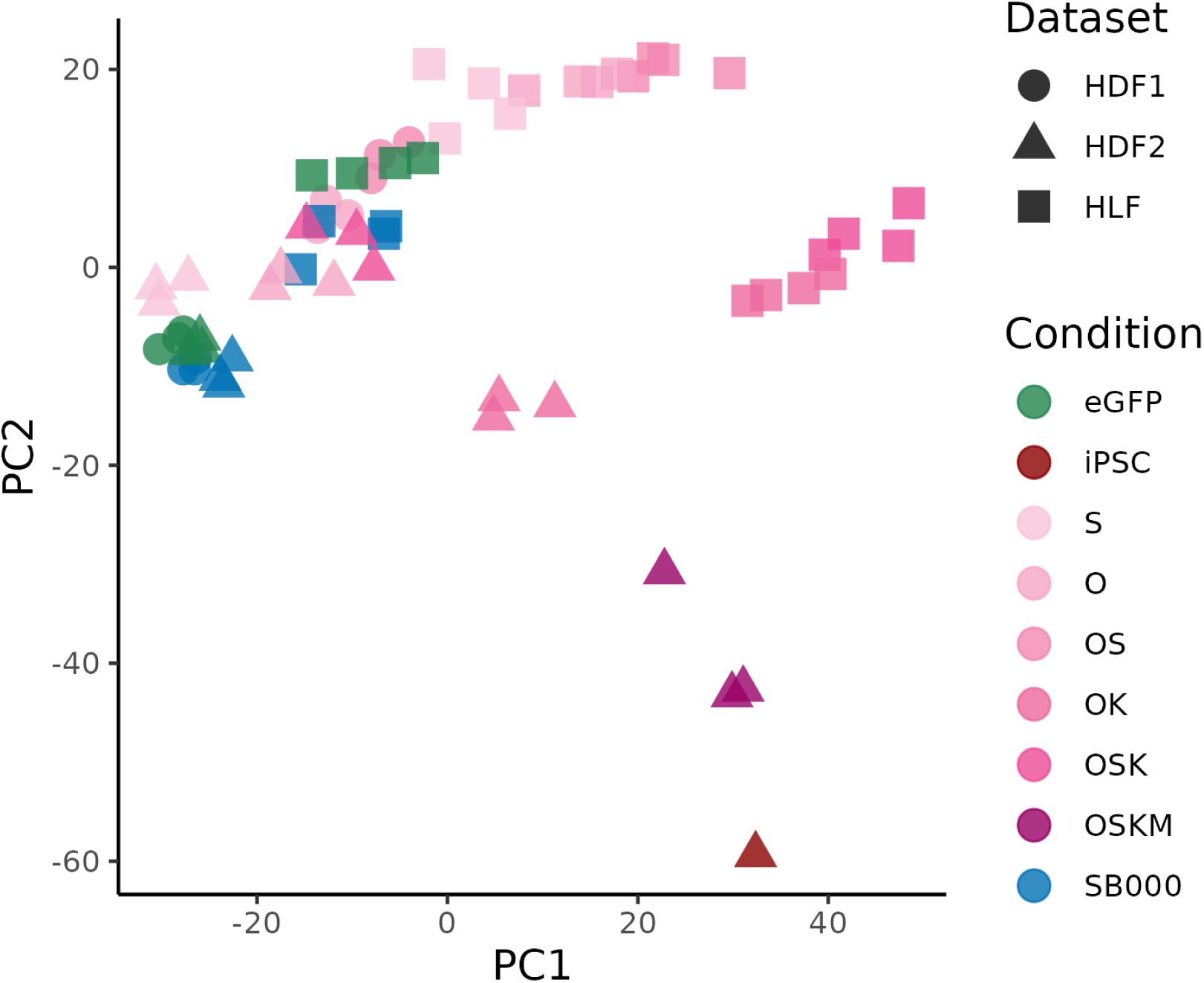
SB000 and the individual Yamanaka factors O, S and K were encoded in lentiviral vectors linking expression of the genes to eGFP. Human dermal fibroblasts (HDFs) and human lung fibroblasts (HLFs), both from aged donors, were exposed to lentiviruses expressing these vectors, eGFP-only vector, no virus or sendai-virus encoded OSKM for two weeks. The transcriptome of cells from each sample was psuedo-bulked and the dimensionality of the data reduced with principal components anaylsis (PCA) after batch correction. Each point represents a sample and is coloured by the intervention it received, with the shape of the point denoting the source of the fibroblasts.

**Figure S7:**
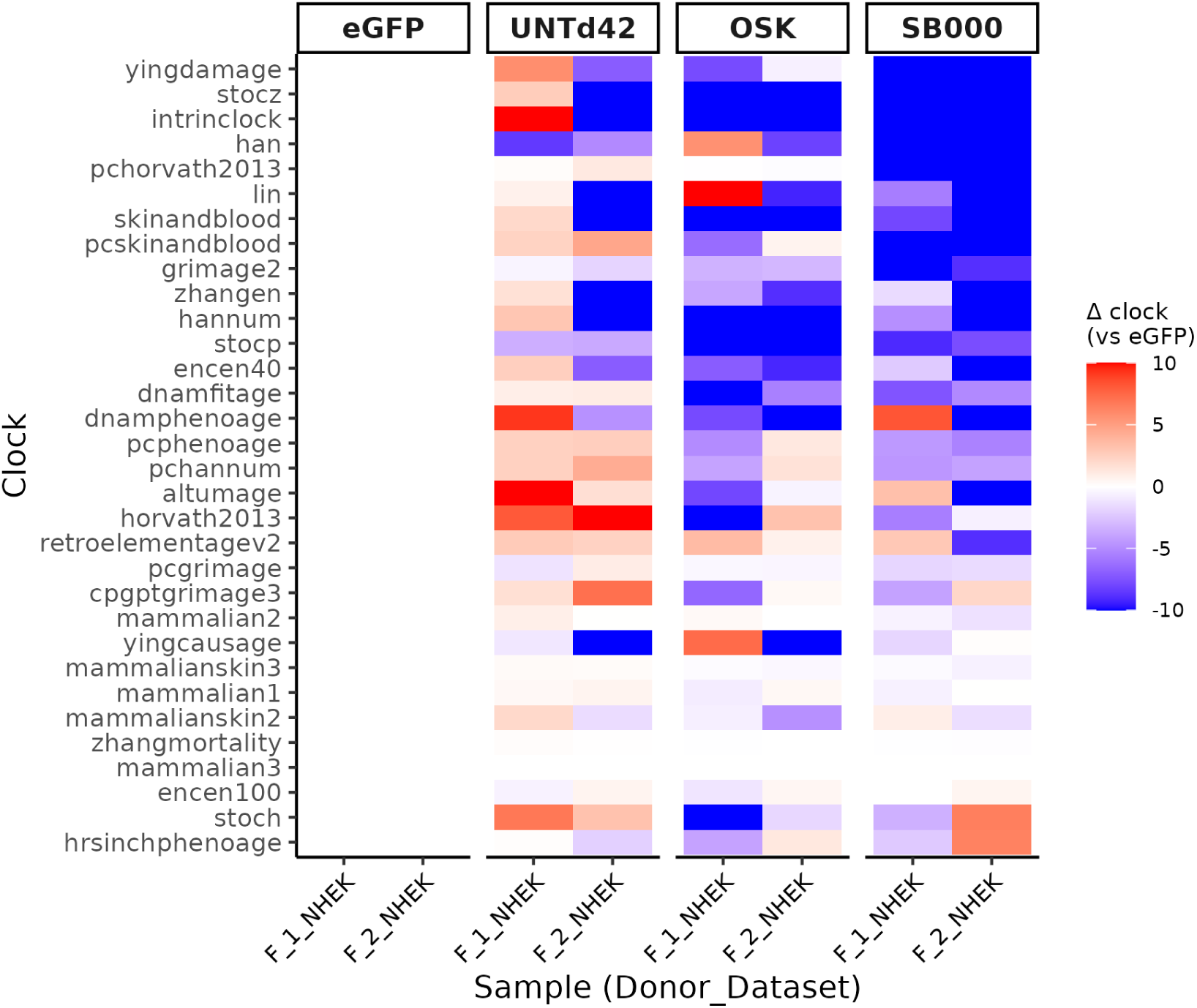
Normal human epidermal keratinocytes (NHEKs) were transduced with SB000, OSK or eGFP control lentivirus, and the fluorescence population sorted using FACS. Six weeks post-transduction, the epigenome of bulk samples was profiled using the EPICv2 array. The epigenetic age of samples was calculated with many published epigenetic clock algorithms using the *pyaging* package; the heatmap shows the change in predicted age of cell samples receiving SB000, OSK or no vector (“UNTd42”), relative to eGFP-receiving cells from the same donor.

**Figure S8:**
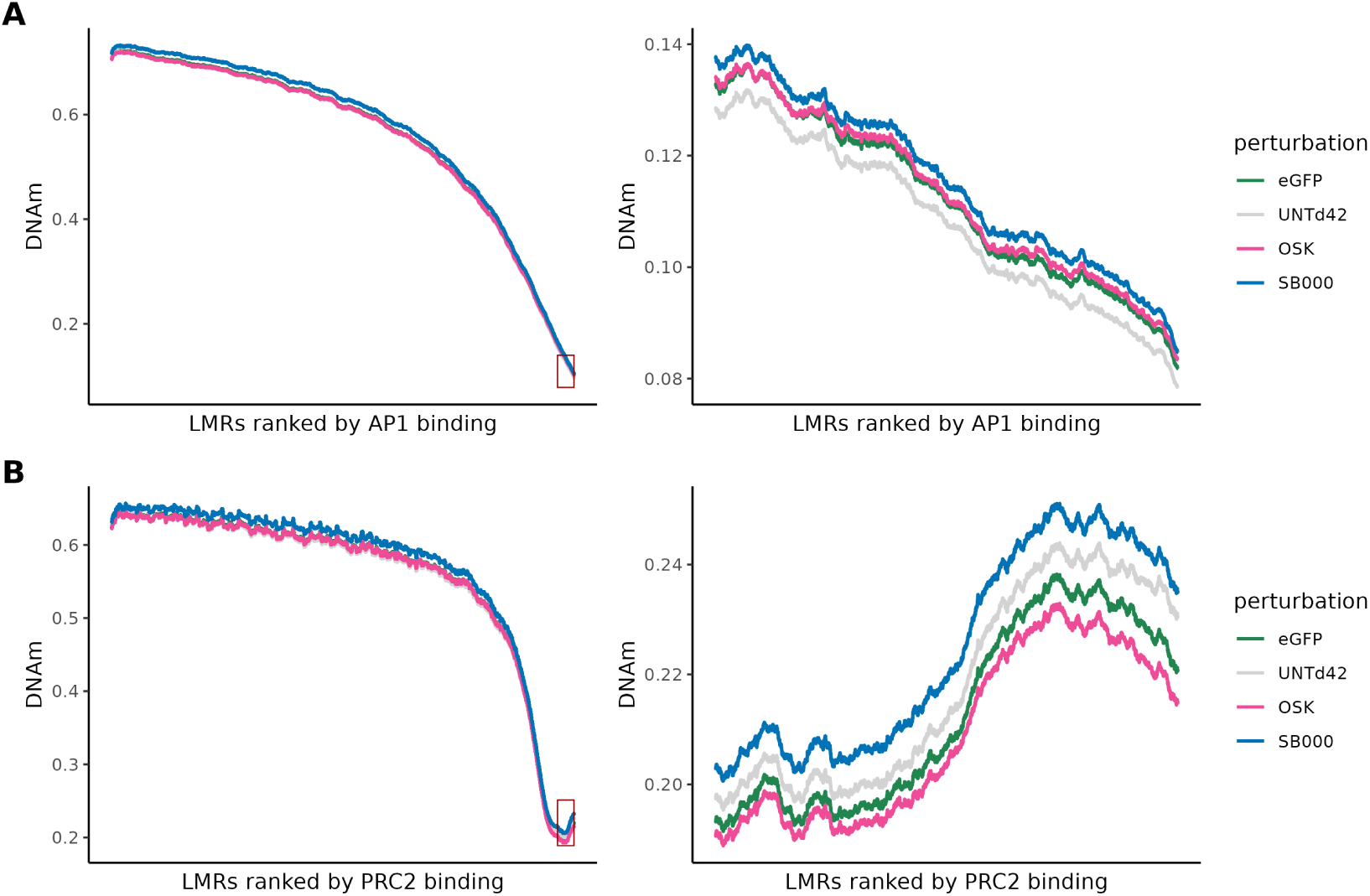
Normal human epidermal keratinocytes (NHEKs) were transduced with SB000, OSK or eGFP control lentivirus, and the fluorescence population sorted using FACS. Six weeks post-transduction, the epigenome of bulk samples was profiled using the EPICv2 array. **(A)**, left: the methylation status of DNA in AP1 binding sites (low-methylated regions, LMRs) across the methylome. The DNAm level at CpG sites is plotted in ascending order of AP1 binding enrichment, with each of the conditions coloured. Right: a zoomed representation of the top 1000 sites most enriched for AP1 binding (highlighted by a red box in the left panel). **(B)**, left: the methylation status of DNA in PRC2 binding sites (LMRs) across the methylome. The DNAm level at CpG sites is plotted in ascending order of PRC2 binding enrichment, with each of the conditions coloured. Right: a zoomed representation of the top 1000 sites most enriched for PRC2 binding (highlighted by a red box in the left panel).

**Figure S9:**
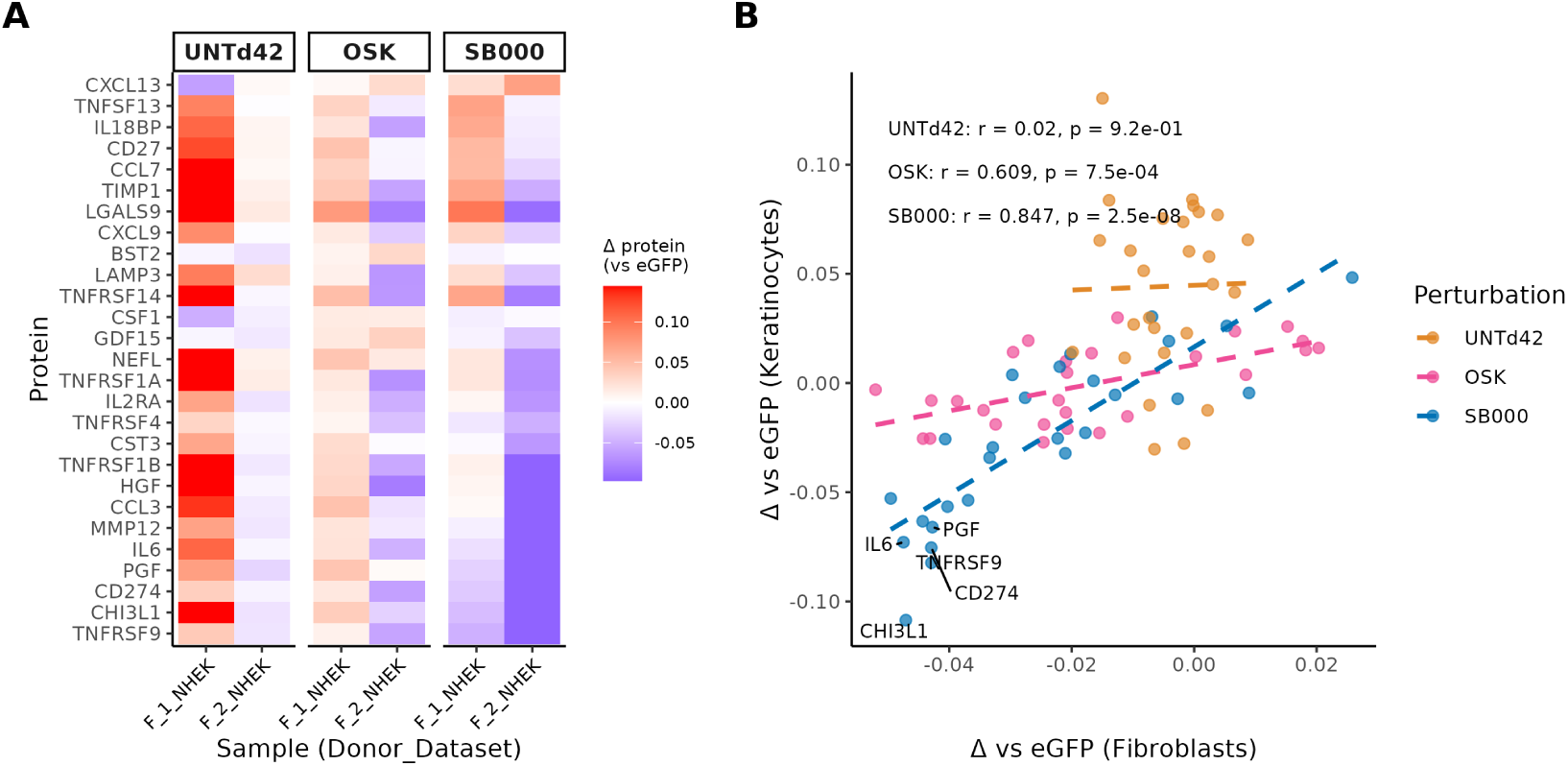
Normal human epidermal keratinocytes (NHEKs) were transduced with SB000, OSK or eGFP control lentivirus, and the fluorescence population sorted using FACS. Six weeks post-transduction, the epigenome of bulk samples was profiled using the EPICv2 array. CpGPT was used to predict the level of circulating mortality-associated proteins from the DNAm profiles of these samples. **(A)**: a heatmap showing the predicted levels of disease-associated blood proteins for untreated cells (“UNTd42”), or those expressing OSK or SB000. Values are expressed relative to those for eGFP samples from the same donor. **(B)**: The mean values for each NHEK condition from **(A)** are plotted on the y-axis, versus the matched condition in HDFs (from Figure S4). A line of best fit for the paired predictions is shown with a dotted line, and the Pearson correlation coefficient for each perturbation is inset. Points and lines are coloured by perturbation.

**Figure S10:**
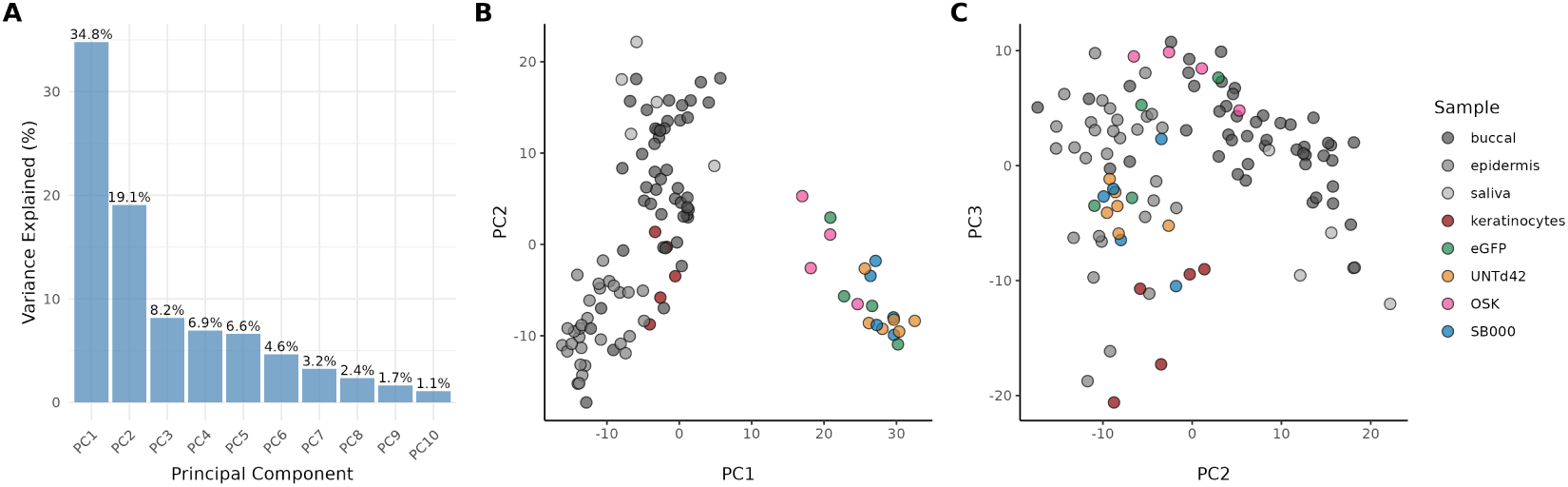
Normal human epidermal keratinocytes (NHEKs) were transduced with SB000, OSK or eGFP control lentivirus, and the fluorescence population sorted using FACS. Six weeks post-transduction, the epigenome of bulk samples was profiled using the EPICv2 array. The DNAm profiles of SB000-, OSK-, and eGFP treated NHEK samples were projected into the embedding space of CpGPT, along with a large reference dataset. To refine the tissue/cell-type similarities to each of our perturbations, we first performed hierarchical clustering on the top 20 PCs from the embeddings of the combined dataset. Then, we performed principal components analysis (PCA) using the subset of samples that were within the same cluster that our data was part of. **(A)** shows the explained variance for each of the principal components from the PCA of this cluster. **(B)** Shows the local cluster in the PC1 and PC2 dimensions, and **(C)** shows the PC2 and PC3 dimensions.

